# Piconewton forces mediate GAIN domain dissociation of the latrophilin-3 adhesion GPCR

**DOI:** 10.1101/2023.01.12.523854

**Authors:** Brian L. Zhong, Christina E. Lee, Vipul T. Vachharajani, Thomas C. Südhof, Alexander R. Dunn

## Abstract

Latrophilins are adhesion G-protein coupled receptors (aGPCRs) that control excitatory synapse formation. aGPCRs, including latrophilins, are autoproteolytically cleaved at their GPCR-Autoproteolysis Inducing (GAIN) domain, but the two resulting fragments remain associated on the cell surface. It is thought that force-mediated dissociation of the fragments exposes a peptide that activates G-protein signaling of aGPCRs, but whether GAIN domain dissociation can occur on biologically relevant timescales and at physiological forces is unknown. Here, we show using magnetic tweezers that physiological forces dramatically accelerate the dissociation of the latrophilin-3 GAIN domain. Forces in the 1-10 pN range were sufficient to dissociate the GAIN domain on a seconds-to-minutes timescale, and the GAIN domain fragments reversibly reassociated after dissociation. Thus, mechanical force may be a key driver of latrophilin signaling during synapse formation, suggesting a physiological mechanism by which aGPCRs may mediate mechanically-induced signal transduction.

## Introduction

Latrophilins are adhesion G-protein coupled receptors (aGPCRs) that have been implicated in multiple neuropsychiatric diseases^1,2^. Postsynaptic latrophilins are highly expressed in the brain and are necessary for excitatory synapse formation in specific brain regions^3-5^. Latrophilin knockout mice exhibit impairments in spatial memory and learning^2^. The mechanism of latrophilin-mediated synapse formation involves coincident binding of two presynaptic ligands, teneurins (Ten) and fibronectin leucine-rich repeat transmembrane proteins (FLRT)^4,6^. How trans-synaptic binding of these ligands to latrophilins induces excitatory synapse formation is unknown, though G-protein activity downstream of latrophilins seems to be necessary^7^.

At the molecular level, latrophilins—like most aGPCRs—are autoproteolytically cleaved at their GPCR-Autoproteolysis Inducing (GAIN) domain that is conserved across the aGPCR family of proteins^8^ (**Fig. 1A**). The resulting latrophilin fragments remain associated on the cell surface^8,9^. The prevailing “tethered agonist” model of aGPCR activation posits that, after GAIN domain mediated autoproteolysis, dissociation of the extracellular N-terminal fragment (NTF) from the transmembrane C-terminal fragment (CTF) liberates an agonist peptide (termed *Stachel*) at the very N-terminus of the CTF. The subsequent binding of this tethered agonist *Stachel* peptide to the 7-helix transmembrane domain (7-TMD) is hypothesized to trigger downstream signaling^10^. Dissociation has been theorized to occur in response to mechanical forces applied to latrophilin through binding of teneurins and/or FLRTs^10,11^, which may transmit forces to latrophilin generated by cytoskeletal processes or relative cell motion. However, mechanically-induced dissociation of the latrophilin NTF and CTF has not been demonstrated, and the necessity of GAIN-mediated autoproteolysis for signaling via aGPCRs more broadly is unclear. In some cases, autoproteolysis appears to be necessary for activation of aGPCRs by their ligands and other small molecule agonists, as has been shown for the GPR56 receptor^12^. Moreover, aGPCR constructs lacking the NTF can trigger greater G-protein activation compared to their full-length counterparts^10,13^, consistent with a model of dissociation-triggered activation. However, non-cleavable latrophilin mutants have been shown to rescue at least some phenotypes of latrophilin knockouts^4,7,14,15^, suggesting that autoproteolysis and dissociation may be dispensable for certain latrophilin functions, including G-protein regulated functions. Thus, while it is possible aGPCR dissociation regulates downstream signaling, whether and how this process could occur physiologically is not well established.

**Figure 1.**
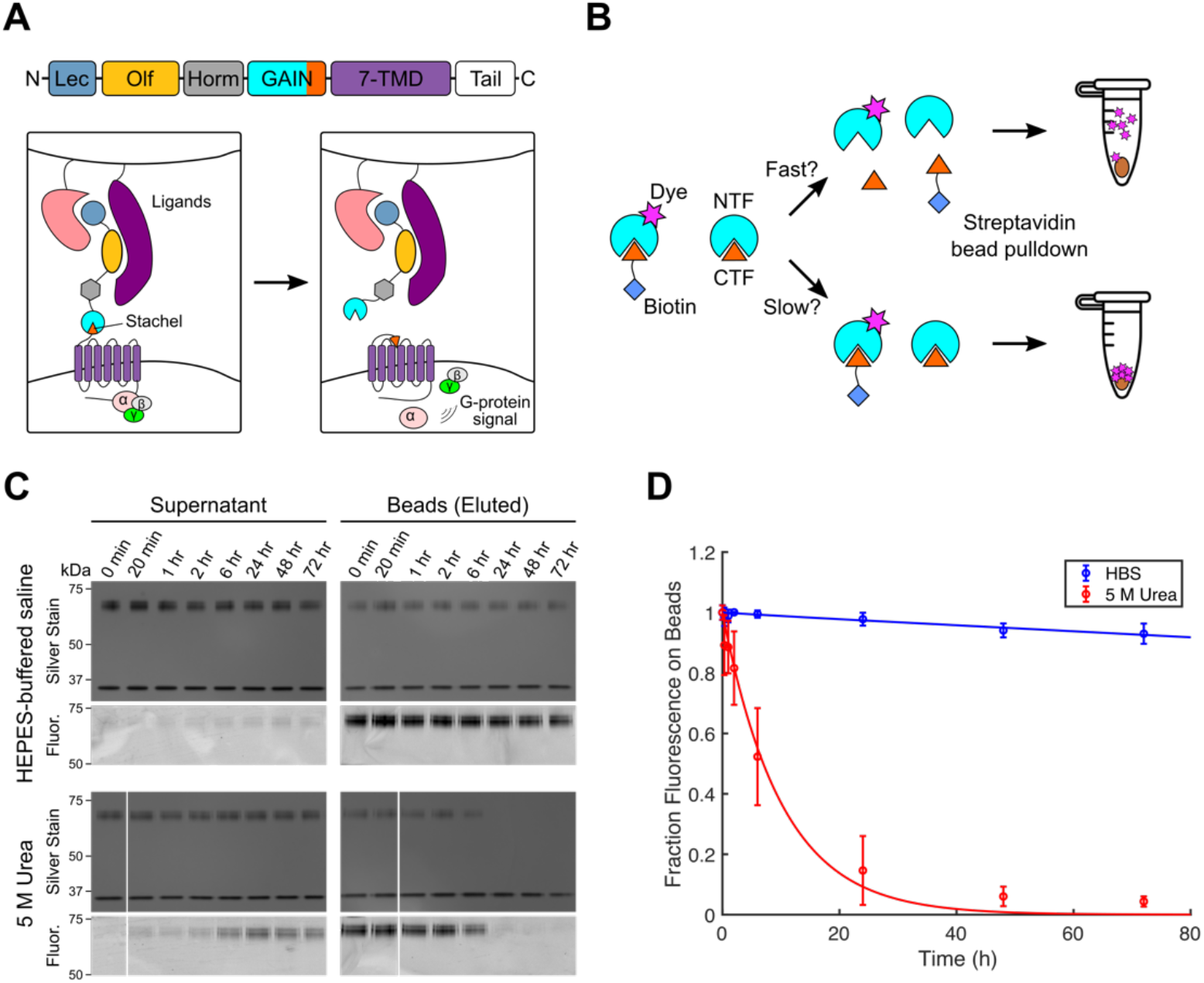
The latrophilin-3 GAIN domain forms a highly stable autoproteolyzed complex that remains associated over the course of days. **(A)** Domain structure of latrophilin aGPCRs and proposed mechanism of signaling mediated by exposure of a tethered agonist peptide (*Stachel*). The lectin (Lec) and olfactomedin-like (Olf) domains are responsible for binding transsynaptic ligands. These domains are followed by a hormone-binding domain (Horm) and the GPCR autoproteolysis-inducing (GAIN) domain where the receptor is cleaved, followed by the 7-pass transmembrane domain (7-TMD) and intracellular tail that regulate G-protein coupling. **(B)** Schematic of streptavidin pulldown assay of labeled GAIN domains to measure dissociation in solution. The N-terminal half of the GAIN domain construct is labeled with a dye, and the C-terminal half is labeled with biotin. The doubly labeled GAIN domain is mixed with excess unlabeled GAIN domain, and the mixture is sampled at various timepoints. Biotinylated protein in each sample is pulled down with superparamagnetic streptavidin beads, and the protein content and fluorescence in both the supernatant and bead fractions are assessed **(C)** Representative silver stained gel of protein in supernatant *(left)* and pulled down on streptavidin beads *(right)* and Cy3 fluorescence at various time points for latrophilin-3 GAIN domain in HEPES-buffered saline *(top)* or 5 M urea *(bottom)*. **(D)** Quantification of the fraction of Cy3 fluorescence in the bead fraction compared relative total fluorescence at various time points in HEPES-buffered saline *(blue)* or 5 M urea *(red)*. Error bars are standard errors for 3 experimental trials. Lines through points are exponential fits with decay constants of 0.0011 hr^-1^ and 0.099 hr^-1^ for Lphn3 in HEPES-buffered saline and 5 M urea, respectively.

Here, we asked whether physiological levels of mechanical force are sufficient to dissociate GAIN domains on timescales commensurate with cell signaling, as suggested by the tethered agonist hypothesis. We used magnetic tweezers assays to measure the effect of mechanical force on the dissociation of the latrophilin-3 (Lphn3) GAIN domain. These experiments reveal that physiological levels of mechanical force in the low piconewton range are sufficient to dissociate the Lphn3 GAIN domain on timescales consistent with cell signaling. These insights illustrate the responsiveness of latrophilins to mechanical force at the molecular level and provide evidence that physiological levels of force are likely to regulate signaling by latrophilin and potentially other aGPCRs.

## Results

### Piconewton forces promote Lphn3 NTF and CTF dissociation on a seconds-to-minutes time scale

Previous studies suggest that the latrophilin NTF and CTF remain associated after autoproteolysis, both when expressed on the cell surface and when isolated as purified protein^8,16^. Consistent with the literature precedent, we isolated a complex of the NTF and CTF after expressing and purifying a secreted Lphn3 GAIN domain protein fused to EGFP at the N terminus and HaloTag at the C terminus (**Supp. Fig. 1**) in mammalian cell suspension culture over the course of 5 days, highlighting the stability of their association. To further quantify the stability of the complex, we used a labeling and affinity pulldown approach. We labeled the Lphn3 GAIN domain fusion protein on the NTF with a Cy3 dye at a YbbR^17^ tag using 4’-phosphopantetheinyl transferase (Sfp synthase) and the CTF with a biotinylated HaloTag ligand. We then mixed this doubly labeled protein with unlabeled GAIN domain protein and pulled down biotinylated CTF at various time points using superparamagnetic streptavidin beads (**Fig. 1B**). Separation between the NTF-CTF complex can be visualized by Cy3 in the supernatant from GAIN NTF that has dissociated from its biotinylated CTF counterpart, and thus, cannot be pulled down. Even after 72 hours in HEPES-buffered saline (HBS), we observed minimal Cy3 fluorescence, and hence NTF, in the supernatant. A majority of the fluorescent protein eluted from the bead fraction (**Fig. 1C, D**), which implies that the stability of the GAIN domain association reflects, at least in part, a remarkably slow off-rate. By contrast, treatment of the GAIN domain with 5 M urea resulted in nearly full dissociation of the GAIN domain 24 hours after starting the treatment (**Fig. 1C, D**). Rate constants extracted from exponential fits of solution dissociation data for Lphn3 are 0.0011 hr^-1^ (95% CI: (0.0009, 0.0013)) in HBS and 0.099 hr^-1^ (95% CI: (0.075, 0.123)) in 5 M urea. These rate constants imply that the Lphn3 GAIN domain fragments are stable over the course of days to weeks in near-physiological buffers, and are surprisingly resistant to dissociation even in a 5 M urea buffer.

Given the stability of the GAIN domain linkage, we next sought to examine whether physiological levels of mechanical force can dissociate the Lphn3 GAIN domain. We used magnetic tweezers to directly apply piconewton (pN) mechanical forces to surface-tethered GAIN domains. We tethered the purified Lphn3 GAIN fusion protein at the C-terminal end to HaloLigand-functionalized glass coverslips and biotinylated the N-terminal fragment of the construct at a YbbR tag using Sfp synthase. We then applied forces ranging from 1-10 pN through superparamagnetic streptavidin beads and measured the number of beads that remained tethered to the coverslip over time (**Fig. 2A**). As expected, surfaces functionalized with biotinylated protein contained greater numbers of beads prior to force application as compared to surfaces functionalized with no protein or non-biotinylated protein (**Supp. Fig. 2A**). We observed dissociation of a substantial fraction of beads when mechanical force was applied to GAIN domain surfaces (**Supp. Fig. 2B**). A majority of beads dissociated within 10-100’s of seconds, with increasing forces inducing faster rates of bead dissociation. Sample traces of normalized bead numbers remaining on Lphn3 GAIN surfaces are plotted in **Fig. 2B**. By comparison, a much smaller fraction of beads dissociated from negative control surfaces covalently coated with cleavage-deficient GAIN domains harboring the T855G mutation^4^ (**Supp. Fig. 2B**). This background level of dissociation likely reflects dissociation of residual beads that are non-specifically adhered to the surface, or potentially rupture of biotin-streptavidin bonds. Moreover, this dissociation occurred at slower rates compared to the bead dissociation on Lphn3 GAIN surfaces (**Supp. Fig. 2B, Supp. Fig. 3**), further supporting the idea that the bead dissociation kinetics observed in **Fig. 2B** are indicative of GAIN domain dissociation kinetics.

**Figure 2.**
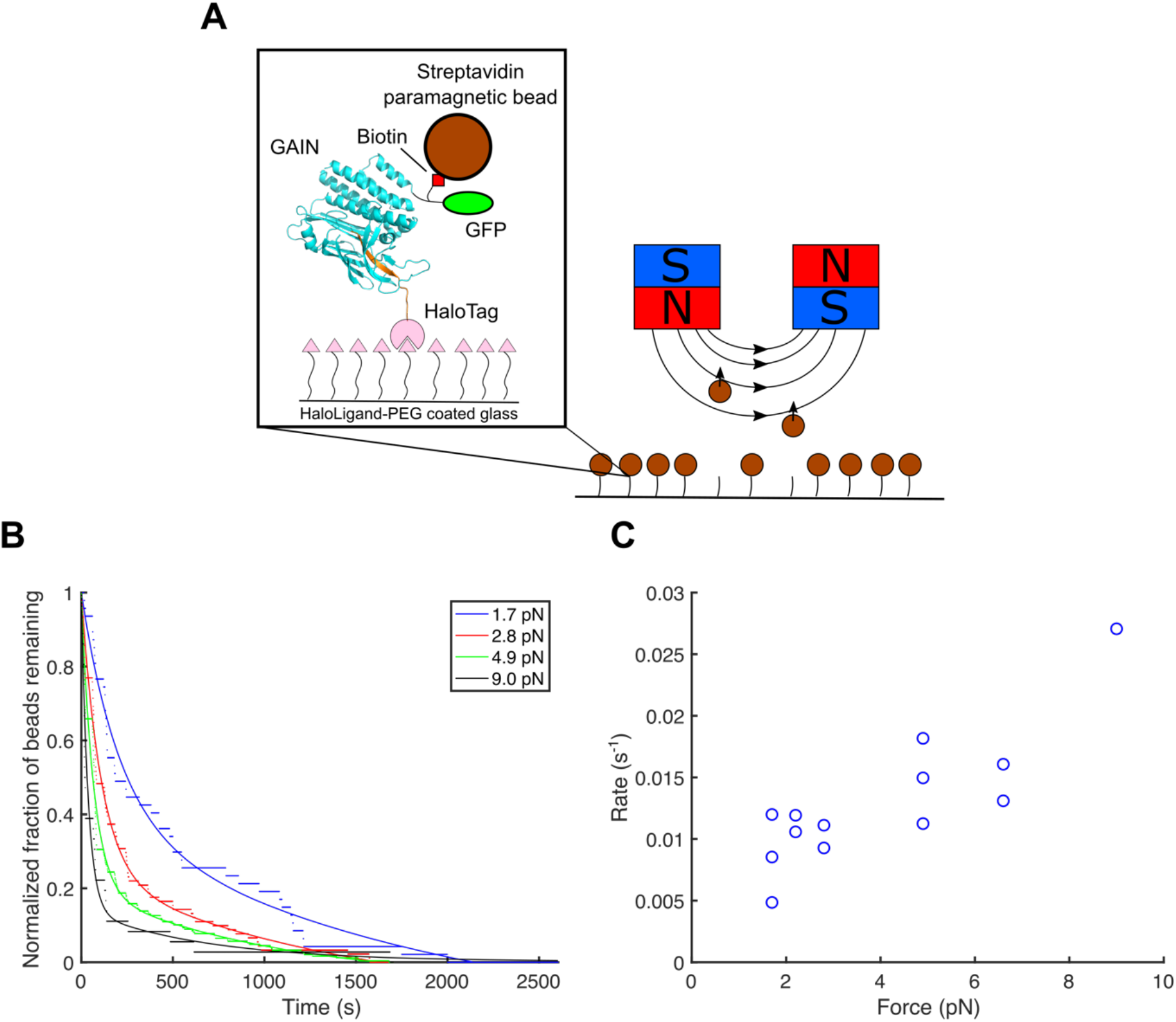
The latrophilin-3 GAIN domain dissociates on the seconds-to-minutes timescale under single-pN forces. **(A)** Schematic of Lphn3 GAIN domain bead dissociation assay and tethering scheme. Molecules of a Lphn3 GAIN domain fusion protein are covalently tethered to a glass surface and functionalized at the other end with biotin for binding with superparamagnetic streptavidin beads. Magnetic tweezers are lowered to a desired height to apply a specific amount of force to the GAIN domain molecules, and the number of beads remaining on the surface is counted over time. **(B)** Representative traces of bead fractions measured over time at 1.7 pN (blue), 2.8 pN (red), 4.9 pN (green), and 9.0 pN (black). Bead fractions computed from measured bead counts are plotted as dots, and curves depict biexponential fits, with a slow rate constant fixed based on rates calculated from dissociation curves of Lphn3 T855G GAIN domains. **(C)** Rate constants of Lphn3 GAIN domain dissociation at various forces, where each point represents one dissociation experiment. Rate constants for the dissociation of the Lphn3 GAIN domain were calculated as the fast rate constant of the biexponential fit to the data.

The lifetime distributions of beads tethered by the Lphn3 GAIN domain were empirically well described by the sum of two exponentials (**Fig. 2B**), suggesting at least two distinct modes of bead dissociation from the surface. The slower of the two dissociation rates measured at a given force was similar to the rate derived from single-exponential fits to the T855G mutant data (**Supp. Fig. 3, Supp. Table 1**), suggesting that these slowly dissociating beads might reflect a distinct subpopulation stemming from a non-specific process independent of GAIN domain dissociation. We therefore fit the Lphn3 GAIN domain bead counts to a biexponential model (**Fig. 2B**), in which the slower rate constant was calculated from T855G measurements at the corresponding force (**Supp. Fig. 3**). In this fit, the faster rate constant represents the rate of dissociation characteristic of the Lphn3 GAIN domain (**Fig. 2C**). In contrast to the high degree of stability observed for the Lphn3 GAIN domain in solution, these data indicate that the bond between the two fragments of the GAIN domain is mechanically labile, with physiological forces reducing the lifetime of the bond by several orders of magnitude.

### A tethered ligand assay demonstrates reversibility of latrophilin-3 GAIN domain dissociation

As an orthogonal strategy to verify our results from the force-mediated bead detachment assay and probe the reversibility of GAIN domain dissociation, we used magnetic tweezers to apply mechanical force to a construct comprising the NTF and CTF of the GAIN domain externally tethered via a GS linker. We expressed the Lphn3 GAIN domain fused to a GS linker with a C-terminal LPETGG tag, which can be covalently ligated onto an N-terminal triglycine using a sortase A mediated reaction to provide a linkage between the two halves, secondary to the non-covalent NTF-CTF interaction (**Fig. 3A**). As such, the two halves of the GAIN domain can remain in close proximity to potentially rebind after dissociation. This “tethered ligand” strategy has been employed to probe the force-dependent dissociation kinetics of other pairs of binding partners^18-21^.

**Figure 3.**
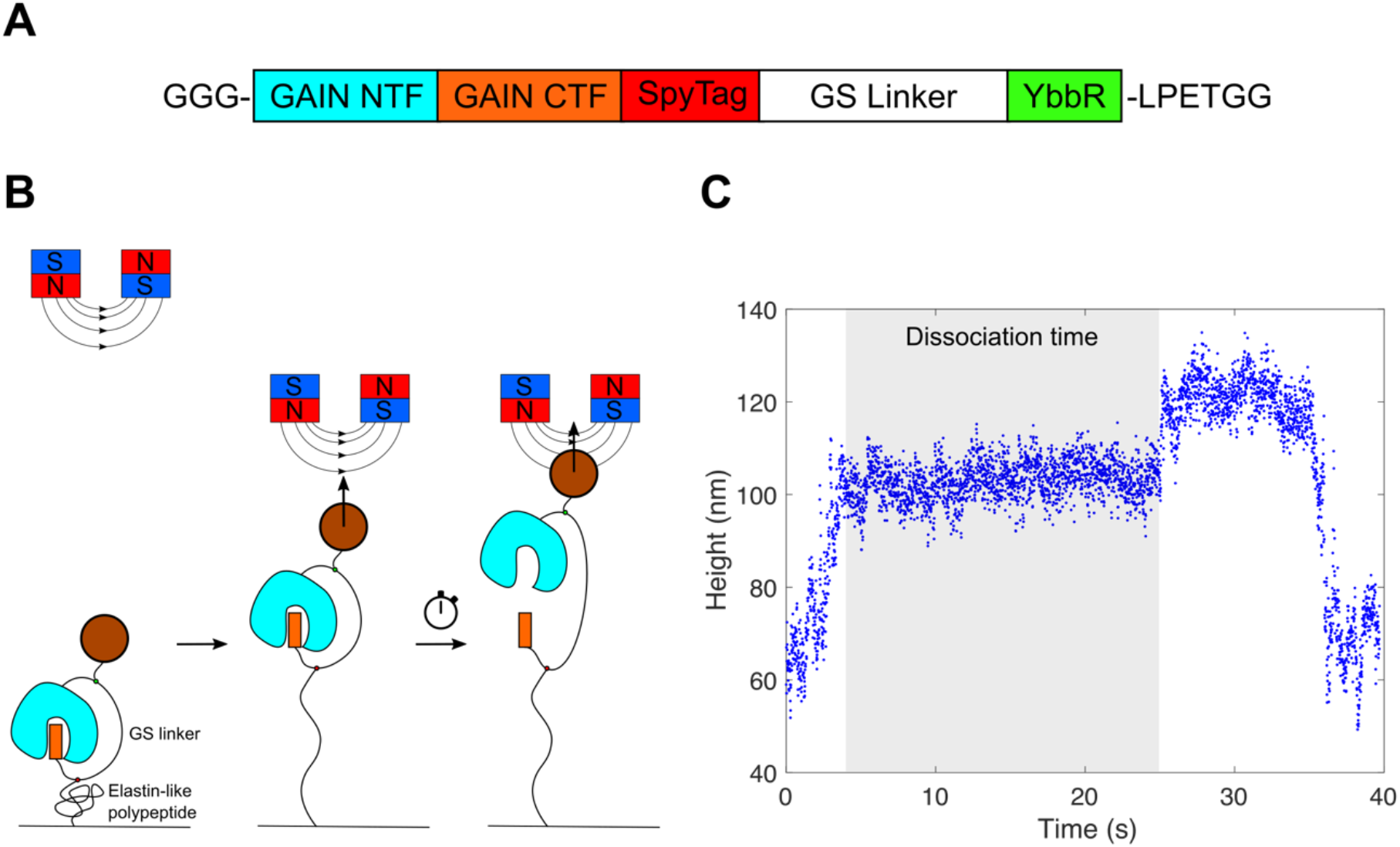
A tethered ligand assay demonstrates reversible dissociation of the latrophilin-3 GAIN domain under single-pN forces. **(A)** Construct design for Lphn3 GAIN domain (after signal peptide cleavage) for tethered ligand assay. C-terminal LPETGG tag can be ligated onto the triglycine motif to synthetically link the NTF and CTF through a GS linker. **(B)** Schematic of force application protocol for the tethered ligand assay. Force is quickly increased from a sub-pN resting state to the desired force, which induces rapid extension of the elastin-like polypeptide. The tethered bead is observed at this force until the height increases again as the GS linker is pulled taut upon NTF and CTF dissociation, and the time between force application and GS linker extension is recorded as the dissociation time (lifetime). **(C)** Sample trace from one round of force application, where the bead height is tracked and the measured dissociation time is highlighted in gray. The force is rapidly increased to the desired force at the start of the trace. The force is then brought back to its low force value after the dissociation of the complex, and the complex is allowed to rebind before the process is repeated.

We used magnetic tweezers to apply piconewton forces to individual molecules of externally ligated, biotinylated Lphn3 GAIN domains through superparamagnetic streptavidin beads and tracked the bead height to determine whether the complex was associated or dissociated (**Fig. 3B**). GAIN domains were covalently bound at an internal SpyTag to a coverslip functionalized with elastin-like polypeptide (ELP) fused to SpyCatcher (**Fig. 3B**). Initial increasing of force from sub-pN to the desired force (typically on the order of 1-10 pN) resulted in rapid extension of the ELP, which served as a fingerprint for specific tethers to GAIN domain complexes (**Fig. 3B, C**). This initial ELP extension was followed by a shorter, secondary extension while the force was held constant, corresponding to the extension of the GS linker due to the dissociation between NTF and CTF of the GAIN domain construct (**Fig. 3C**). The time between the initial force ramp and the dissociation-mediated extension was recorded as the dissociation time. The force was subsequently brought back down to sub-pN levels after dissociation to allow the two halves to rebind in preparation for the next measurement.

Measured dissociation times as a function of force are plotted in **Supp. Fig. 4** and **Fig. 4A**. We measured 131 dissociation events from 19 molecules (**Supp. Fig. 4**), which interestingly implies that the dissociated NTF and CTF can rebind and that dissociation of the Lphn3 GAIN domain is reversible. However, rebinding seems to occur slowly relative to our observed rates of dissociation, and we often observed changes in bead height corresponding to a direct transition to the fully-extended linker height, indicative of force application on a molecule that did not yet rebind (**Supp. Fig. 5**). We estimate based on the observed likelihood of rebinding at various wait times for the tethered ligand construct that the average wait time between rebinding of the NTF and CTF after dissociation is 555 s (**Supp. Note 1**). Thus, while the two halves of the Lphn3 GAIN domain may be able to rebind, this process occurs relatively slowly and may not be physiologically significant for latrophilin signaling. Consistent with the dissociation rates extracted from the bead counting assay, application of single-pN forces to the externally tethered Lphn3 GAIN domains yielded average dissociation times ranging from 10s to 100s of seconds (**Fig. 4A, Supp. Fig. 4**). The agreement of measurements from these two assays suggests that the mechanical behavior observed in these experiments is likely specific to the dynamics of the Lphn3 GAIN domain (**Supp Fig. 4**) and that physiological levels of mechanical force can substantially increase the rate of GAIN domain dissociation.

**Figure 4.**
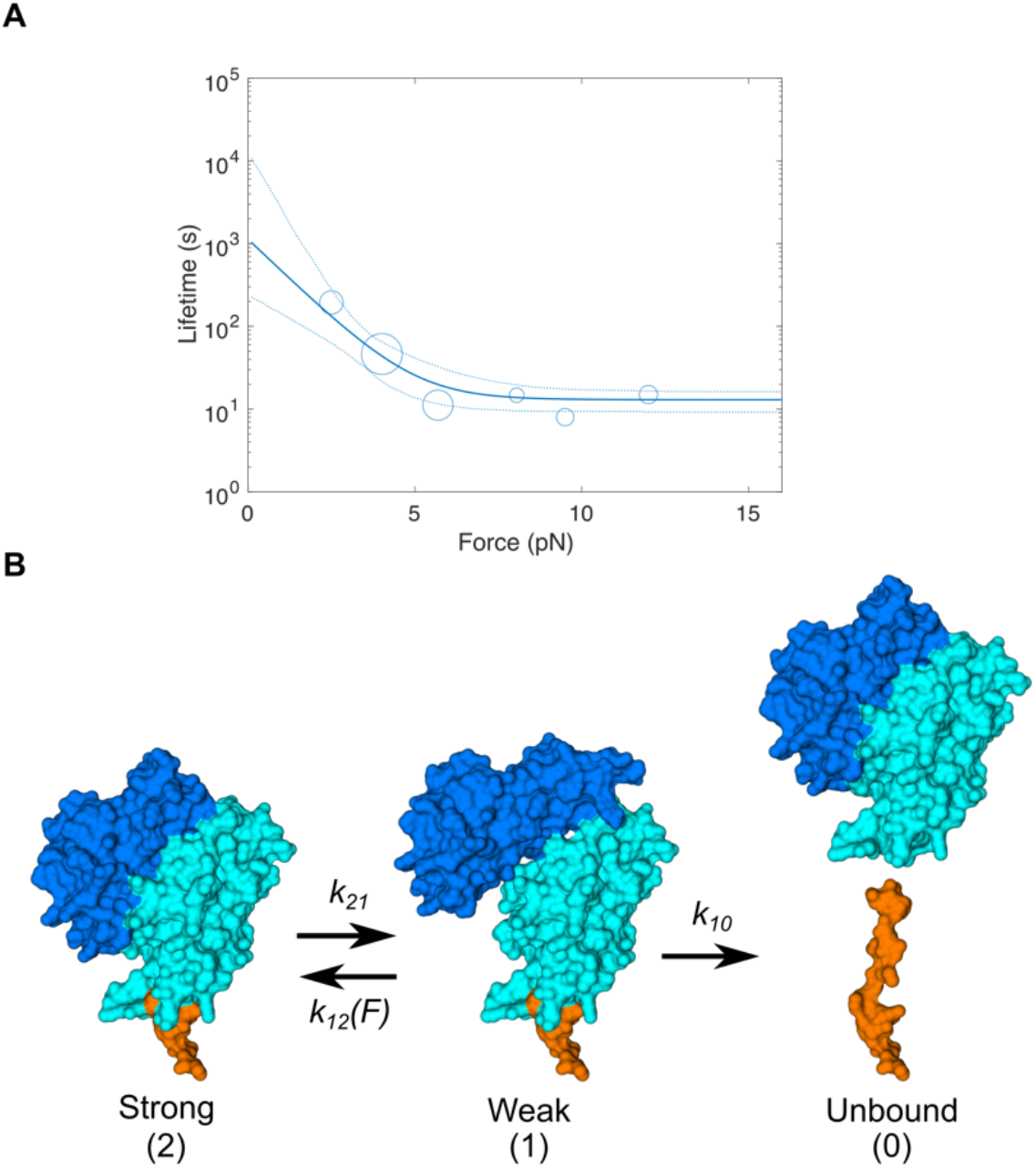
Modeling force-mediated dissociation of the latrophilin-3 GAIN domain. **(A)** Binned lifetime data at measured force ranges from tethered ligand assay (circles) with predicted lifetimes from maximum likelihood estimate fitting to a 2-state dissociation model plotted as a solid blue line. 95% confidence intervals for lifetimes are plotted as dotted blue lines. Circles are average lifetimes from bins with 2 pN width, and sizes of circles correspond to number of points in each bin. Data are binned from 131 dissociation events across 19 molecules **(B)** In a proposed kinetic model, the GAIN domain converts between strongly- and weakly-bound states, and the transition from weakly- to strongly-bound states is force sensitive. Dissociation of the GAIN domain occurs from the weakly-bound state, and the dissociation transition itself is relatively force-insensitive. The weak state may represent a structural rearrangement of the GAIN domain, as illustrated in the cartoon as a hypothetical intermediate. Illustration is based on the crystal structure of the rat latrophilin-1 GAIN domain (PDB: 4DLQ)^8^

### Modeling the mechanism of force-mediated latrophilin dissociation

Our dissociation lifetime data illustrate that the Lphn3 GAIN domain appears to be most mechanically sensitive at forces below ∼6 pN, with bond lifetimes decreasing rapidly from our lowest measured forces at ∼2 pN and plateauing at ∼10 s at forces above 6 pN (**Fig. 4A, Supp. Fig. 4**). We fit these lifetime data using maximum-likelihood estimation to different force-dependent kinetic slip-bond models, in which force accelerates the dissociation of a bound complex. We found that a model with 2 bound states yielded higher likelihood values than a model in which a single bound state unbinds according to Bell-Evans kinetics^22^ (**Supp. Fig. 6**). Specifically, the data are well-described by a 4-parameter model in which the Lphn3 GAIN domain interconverts between weakly and strongly bound states (denoted *1* and *2*, respectively) and transitions to an unbound state (denoted *0*) from the weakly bound state (**Fig. 4B**). The force sensitivity of the complex stems from the force sensitivity of the transition from the weakly bound state (*1*) to the strongly bound state (*2*), given by

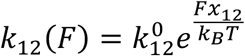

where *k*_*B*_ is Boltzmann’s constant, *T* is temperature, and the distance parameter x_12_ is constrained to be negative so that increasing force favors the weakly-bound state. The interaction between the NTF and CTF in the weak state can be largely modeled as an ideal bond that is insensitive to force. Values for the rate and distance parameters in this model are tabulated in **Table 1**. Rate constants at zero force suggest that the strongly-bound GAIN domain is favored nearly 1000-fold relative to the weakly-bound state from which dissociation occurs, which may help to explain the solution stability of the GAIN domain. The distance parameter *x*_*12*_ represents a distance between the weakly bound GAIN domain and the transition state between the two GAIN domain states. This parameter can be roughly interpreted as a length scale associated with conformational changes between the weakly-bound state and the transition state, and the maximum-likelihood (absolute) value for this parameter of 3.67 nm is consistent with plausible structural changes in the GAIN domain.

**Table 1.** Maximum likelihood estimate parameters for 2 state, 4 parameter model of Lphn3 GAIN domain dissociation.

| Parameter | Maximum likelihood estimate | 95% confidence interval |
| --- | --- | --- |
| $k_{10} (\text{s}^{-1})$ | 0.077 | (0.062, 0.107) |
| $k_{12}^0 (\text{s}^{-1})$ | 0.655 | (0.113, 32.8) |
| $x_{12} (\text{nm})$ | -3.67 | (-6.82, -2.07) |
| $k_{21} (\text{s}^{-1})$ | 0.00761 | (0.0037, 0.0522) |

The predicted mean dissociation times for the Lphn3 GAIN domain from the model are plotted in **Fig. 4A**, along with the binned measured dissociation times of the tethered Lphn3 GAIN construct. The Lphn3 GAIN domain bond lifetimes predicted by this model also fit the biexponential trend of lifetimes within specific force bins (**Supp. Fig. 7**), further suggesting that the 2-state model is preferred to the 1-state model. Interestingly, the model predicts an extrapolated lifetime of ∼1000 s at 0 pN force, which is substantially shorter than what is observed for the GAIN domain in solution. Thus, force may dissociate the Lphn3 GAIN domain through pathways distinct from any dissociation that may occur in the absence of force.

## Discussion

Here, we demonstrate using two magnetic tweezers assays that low pN forces are sufficient to dissociate the Lphn3 GAIN domain on a seconds-to-minutes timescale. Previous studies have suggested that aGPCR signaling can be activated by GAIN domain dissociation or treatment with synthetic tethered agonist peptides over the course of minutes^16,23,24^. While these studies did not specifically test the effects of force-mediated dissociation on aGPCR signaling, the dissociation times for the Lphn3 GAIN domain measured in this study are consistent with the timescales reported in the literature for signaling via latrophilins and other aGPCRs. Previous studies on latrophilins and other aGPCRs have established a link between force and signaling^25,26^, but the work presented here represents to the first study to our knowledge that characterizes forces involved in aGPCR dissociation at the molecular level. In the context of synaptogenesis, it remains to be tested whether bonds between latrophilins and ligands such as teneurins or FLRTs can support the forces necessary to dissociate latrophilins at the timescales that we observe. However, the range of forces that we test in this study are relatively modest and accessible compared to forces borne and transmitted by other cytoskeletal proteins and receptors^27-33^. Additionally, based on the solution dissociation kinetics of the GAIN domain and the reassociation rates estimated from the tethered ligand assay, we estimate the binding affinity (K_d_) of the NTF-CTF interaction to be 2.8 μM (see **Supp. Note S1**). This affinity is lower than reported binding affinities for teneurins and FLRTs to latrophilins—measured to be on the order of 10-100s of nM^6,34^—implying that these transsynaptic ligand interactions may be stronger than the intramolecular interaction between latrophilin fragments. Thus, it seems likely that physiological forces at synapses mediated by transsynaptic binding can promote the dissociation of and signaling through latrophilins. These data thereby support the current dominant model of aGPCR signaling that proposes that GAIN domain dissociation can enable a tethered agonist to activate the receptor.

While our results suggest that the Lphn3 GAIN domain can dissociate on relatively short timescales under physiological force, these data do not preclude the possibility that other conformational changes or molecular events independent of dissociation may regulate latrophilin signaling. On the contrary, our data are consistent with a model in which the force sensitivity of GAIN domain dissociation is mediated by a transition between strongly and weakly bound states that precedes dissociation of the complex. It is entirely possible that non-cleavable variants of Lphn3 and other aGPCRs may signal via transitions between these states or through other dynamic conformational changes. Although crystal structures of the latrophilin and other aGPCR GAIN domains suggest a stereotyped secondary structure^8^, recent cryo-EM data on several aGPCRs observed no high-resolution densities for the GAIN domain N-terminal to the tethered agonist^16,35^. These findings may imply that the GAIN domain itself is structurally dynamic. In the context of signaling mediated by the tethered agonist, molecular dynamics studies have suggested that conformational fluctuations in the absence of force may be sufficient to expose parts of the tethered agonist peptide^36^. The force sensitivity of the Lphn3 GAIN domain at low forces may accelerate these transitions to promote tethered agonist exposure, even without full dissociation of the complex. Additionally, several members of the aGPCR family are not observed to be cleaved despite the presence of a GAIN domain^8,13,37^, and increasing evidence has suggested that key residues of the tethered agonist can bind the transmembrane region even without autoproteolysis of the receptor^16,35,36^. Could dissociation-dependent and dissociation-independent modes of force-mediated signaling provide different cues to downstream G-protein signaling pathways? More broadly, it is not presently known whether dissociation of the latrophilin GAIN domain promotes or inhibits synapse formation. Further cellular studies to dissect the contributions of force and dissociation on downstream signaling pathways will be required to provide a clear answer to these questions.

Our measurements of the Lphn3 GAIN domain dissociation rate in solution suggest that the bond between GAIN domain fragments is extremely stable with minimal observed dissociation even over the course of days. This stability is further highlighted by the observation that the GAIN domain only dissociates in hours even when exposed to harsh chemical denaturants, such as 5 M urea. By contrast, extrapolation of our model of force-mediated dissociation to zero force suggests a dissociation lifetime on the order of 1000s of seconds, which is substantially shorter than what is observed for the GAIN domain in solution. It is possible that force-mediated dissociation occurs through pathways different from what would be predicted in the absence of force, or alternatively that additional force-sensitive states not detected in our experiments modulate the overall rate of dissociation at forces too low to be probed in our assay.

Strikingly, our results also suggest that the N- and C-terminal fragments of the Lphn3 GAIN domain may be able to reversibly reassociate after dissociation. However, it is unclear whether such behavior occurs in a cellular context or plays a role in signaling. While some previous studies have suggested that latrophilin fragments may act as independent fragments capable of reassociating^38^, a majority of studies argue for stable association of the fragments after proteolysis and against the notion that the fragments can dissociate and reassociate^4,8,24^. Perhaps relatedly, it is possible that the GAIN-domain dissociation of aGPCRs does not serve to activate signaling, but instead enables internalization of aGPCRs, possibly via β-arrestin signaling^24,39^. Such a role of the GAIN-domain cleavage of aGPCRs is potentially attractive because it would suggest a regulation of the lifetime of aGPCRs on the cell surface by ligand-induced receptor dissociation. Internalization would compete with any rebinding-mediated signaling, particularly since reassociation of the fragments appears to occur slowly relative to the dissociation times under the modest forces probed in this study.

Our results suggest that force may be a critical regulator of latrophilin and aGPCR signaling (either by aGPCR activation or inactivation), but our findings characterize only one domain of one specific aGPCR. Interactions between other domains, particularly between extracellular domains and residues on the transmembrane region, may also play a role in the force sensitivity and dissociation of latrophilins and aGPCRs in a cellular setting. Other cleavage sites, which have been reported for latrophilins^40,41^, may further complicate the interplay between force and dissociation. Moreover, it is possible that other aGPCRs may be sensitive to different forces than those reported here as the GAIN domains of some other aGPCRs may not be as stably associated as latrophilins^16,42^. Nevertheless, our measurements provide a benchmark for the forces necessary to dissociate the GAIN domain of an aGPCR, which is likely to play a role in regulating the force sensitivity and dissociation of the receptor. Our model describing these results also suggests a conceptual framework for the mechanism of force-responsiveness in latrophilins and potentially other aGPCRs. Interestingly, recent functional studies on the GAIN domain of the aGPCR ADGRE5 has highlighted the importance of an α-helical bundle in the function of the GAIN domain as it pertains to cellular signaling^43^. This may hint at a mechanism by which the helical bundle regulates the force-sensitivity and stability of the GAIN domain. Further studies examining the role of force on aGPCR signaling in cellular systems will be necessary to elucidate these details of these mechanisms. Nevertheless, our observations demonstrate that physiological levels of mechanical force are likely to play a key role in regulating signaling by latrophilins and other aGPCRs.

## ACKNOWLEDGEMENTS

B.L.Z is supported by the Stanford ChEM-H Chemistry/Biology Interface Predoctoral Training Program (NIH 5T32GM120007) and the NSF Graduate Research Fellowship Program (DGE-1656518). C.E.L. is supported by the Stanford Molecular Biophysics Training Program (NIH T32GM136568). V.T.V. is supported by the Stanford Medical Scientist Training Program (NIH T32GM007365) and the NIH NIDDK (1F30DK124985). T.C.S acknowledges the NIMH (MH126929), and A.R.D. acknowledges the HHMI (Faculty Scholar Award) and the NIH (R35GM130332). We would also like to acknowledge the Stanford Wu Tsai Neurosciences Institute for providing a seed grant for this project. We thank Dr. Jacob Mahoney, Elise Bruguera, and Professor William Weis for protein expression advice and resources and Robert Lee and Professor Jennifer Cochran for protein purification advice and resources. We are grateful to Dr. Magnus Bauer and Dr. Daniel Matus for discussion and comments on the manuscript.

## AUTHOR CONTRIBUTIONS

B.L.Z., T.C.S., and A.R.D. conceived the study and designed experiments. B.L.Z. and C.E.L. conducted bead dissociation assays and data analysis. B.L.Z. and V.T.V. conducted tethered ligand magnetic tweezers assays. B.L.Z. performed all other experiments, procedures, and data analysis. B.L.Z. and A.R.D. wrote and prepared the manuscript, with input from all other authors.

## DECLARATION OF INTERESTS

The authors declare no competing interests.

## Materials and Methods

### DNA constructs

Solution dissociation and bead dissociation assays were performed using a construct comprised of an N-terminal Ig-κ secretion signal sequence followed by EGFP, a YbbR tag^17^, the *H. sapiens* Lphn3 GAIN domain up to the transmembrane region (Uniprot: Q9HAR2-2, amino acids 551-878), HaloTag^44^, and a human rhinovirus (HRV) 3C protease site cloned into a pcDNA3 vector containing human IgG Fc derived from pcDNA3-Nrxn1beta AS4(-)-Fc (a gift from Peter Scheiffele & Tito Serafini, Addgene plasmid # 59313 ; http://n2t.net/addgene:59313 ; RRID:Addgene_59313). The C-terminal Fc tag was included to facilitate purification via a protein A affinity column. This construct was assembled by Epoch Life Sciences Inc. (Missouri City, TX). A non-cleavable T855G mutant of this Lphn3 GAIN domain fusion protein was generated using QuikChange II XL (Agilent) site-directed mutagenesis with primers 5’-ATG CAG CTG TAA TCA CCT GGG CAA CTT TGC TGT CCT GAT G -3’ and 5’-CAT CAG GAC AGC AAA GTT GCC CAG GTG ATT ACA GCT GCA T -3’.

For tethered ligand assays, the *H. sapiens* Lphn3 GAIN domain was flanked at the N terminus by an Ig-κ secretion signal sequence immediately followed by GGG and at the C terminus by SpyTag followed by a 144 amino acid GS linker, a 6xHis tag, a HRV 3C protease site, a YbbR tag, and a LPETGG sortase A tag^45^. The GS linker was comprised of 4 shorter linker segments made up of alternating SEG and GSAT linkers available from the iGEM parts database (accession numbers BBa_K404300, BBa_K243029)^18^. This construct was assembled by Epoch Life Sciences Inc. A bacterial expression construct (Cys-ELP-SpyCatcher) encoding an N-terminal cysteine, an elastin-like polypeptide of contour length 120 nm fused to a C-terminal minimal SpyCatcher protein and a 6xHis tag was a kind gift from Hermann Gaub. This construct is analogous to the ELP construct reported in Ott *et al*.^46^ but with a C-terminal minimal SpyCatcher and 6xHis tag in place of a sortase A tag. The sequence of the protein encoded by this plasmid is as follows:

MC-[(VPGEG)-(VPGVG)_4_-(VPGAG)_2_-(VPGGG)_2_-(VPGEG)]_6_-LPEDSATHIKFSKRDEDGKELAGATMELRDSSGKTISTWISDGQVKDFYLYPGKYTFVETAAPDGYEVATAITFTVNEQGQVTVNGHHHHHH

A plasmid encoding the phosphopantetheinyl transferase Sfp from *B. subtilis* (Sfp synthase) was a kind gift from Dr. Michael Burkart (Addgene plasmid # 75015 ; http://n2t.net/addgene:75015 ; RRID: Addgene_7015).

### Protein expression and purification

Cys-ELP-SpyCatcher was expressed in NiCo21 (DE3) *E. coli*, which were induced at an optical density (OD) of 0.6 with 1 mM IPTG and incubated at 37 ºC for 6 hours to express protein. Cells were pelleted and resuspended in 50 mM Tris (pH 7) supplemented with 500 μg/mL lysozyme and 1x cOmplete EDTA-free protease inhibitor cocktail (Roche, 11873580001). The pellet was incubated at 4 ºC while rotating end over end for 45 min.

Dithiothreitol (DTT) was added to the cell suspension at 1 mM and the cell suspension was subject to sonication to lyse the bacteria. The sonicated suspension was spun at 13,000×*g* for 30 min at 4 ºC, and the supernatant from this centrifugation step was spun at 13,000×*g* for another 15 min. The supernatant from the second spin was then subject to 2 cycles of a thermoprecipitation and redissolution purification strategy as described previously^46^, collapsing the ELP in 1 M acetate (pH 2.5) and redissolving in 50 mM Tris + 1 mM DTT (pH 7). The purified ELP was snap frozen and stored in 50 mM Tris + 1 mM DTT (pH 7) at -80 ºC until use.

Lphn3 GAIN domain constructs were expressed in Expi293 human embryonic kidney cells (Thermo Fisher, A14527) in suspension culture in Expi293 media (Thermo Fisher, A1435101) at 37 ºC and 8% CO_2._ 20-50 mL of Expi293 culture was seeded at 3×10^6^ cells/mL. 6 hours later, cells were transfected with 1.5 μg DNA per mL of culture and 2.33 μL of Expifectamine (Thermo Fisher, A14524) transfection reagent per μg of DNA in OptiMEM. Enhancers were added according to manufacturers’ instructions 24 hours after addition of DNA and Expifectamine. Expi293 cells were pelleted 4 days after initial transfection, and the supernatant was collected for affinity purification.

For Fc-tagged Lphn3 GAIN constructs, the collected supernatant was diluted 2-fold into HEPES-buffered saline (10 mM HEPES, 150 mM NaCl, pH 7.4, hereafter referred to as HBS) then loaded into a column packed with 500 μL of Pierce Protein A agarose (Thermo Fisher, 20334). The diluted supernatant was allowed to drain by gravity. The column was then washed with 3 × 5 mL HBS. To cleave the Fc tag off of the bound protein, 110 μL of 2.5 μM 6xHis-tagged HRV 3C protease was flowed into the column bed, the bottom of the column was capped, and then a 110 μL HBS stacker was added and the cleavage was allowed to proceed overnight at 4 ºC to give EGFP-YbbR-Lphn3 GAIN-HaloTag. The cleaved protein was then eluted with 3 × 1 mL HBS, and the 3C protease was removed using Dynabeads magnetic beads designed for His-tagged protein isolation (Thermo Fisher, 10103D). The supernatant from the magnetic bead pulldown was then concentrated using Amicon 10 kDa centrifugal filter units (MilliporeSigma), snap frozen, and stored at -80 ºC in HBS until further use.

For 6xHis-tagged tethered ligand assay constructs, the supernatant was first buffer exchanged into a phosphate buffer containing 50 mM sodium phosphate, 300 mM NaCl, and 10 mM imidazole at pH 8 to remove chelating agents in the Expi293 media that could interfere with the binding to nickel. The buffer-exchanged supernatant was loaded into a column containing 1 mL of HisPur Ni-NTA resin slurry (Thermo Fisher, 88222) and allowed to drain by gravity. The column was washed with 3 × 6 mL wash buffer containing 50 mM sodium phosphate, 600 mM NaCl, and 20 mM imidazole at pH 7.4 then eluted with 4 × 1.5 mL elution buffer containing 50 mM sodium phosphate, 300 mM sodium chloride, and 250 mM imidazole at pH 7.4. To externally ligate the NTF and CTF of the tethered ligand Lphn3 construct, 5 μM GAIN domain protein was mixed with 15 μM sortase A in 50 mM Tris, 150 mM NaCl, 10 mM CaCl_2,_ pH 7.5 and reacted overnight at 4 ºC. The reaction mixture was then purified using size-exclusion chromatography in HBS with a Superdex 200 Increase 3.2/300 column (GE Healthcare, 28-9909-46) to remove excess sortase A and undesired reaction byproducts. The fractions containing the desired product were concentrated, snap frozen, and stored at -80 ºC until further use.

### GAIN domain solution dissociation assays

5 μM purified EGFP-YbbR-Lphn3 GAIN-HaloTag protein in HBS was mixed with 5.2 μM HaloTag PEG-Biotin Ligand (Promega, G8592) and incubated at room temperature for 2 hours to biotinylate the CTF of the GAIN construct. 10 mM MgCl_2_, 0.5 μM Sfp synthase, and 4 μM CoA-Cy3 (SiChem, SC-1143) was subsequently added to this GAIN domain reaction mixture to label the NTF of the construct with Cy3 dye. The final concentration of GAIN domain protein in this reaction was 4 μM, and the reaction was allowed to proceed overnight at room temperature.

Prior to the start of the dissociation measurement, Dynabeads M-270 streptavidin superparamagnetic beads (Thermo Fisher, 65305) were prepared as follows: For each condition (e.g., HBS or 5 M urea), 50 μL of beads were washed by mixing with 250 μL of HBS + 0.1% Tween-20. Beads were then collected by pull down with a magnet, and the supernatant was aspirated. Beads were similarly washed twice more with 250 μL of HBS, then finally resuspended in 80 μL HBS and aliquoted into 10 μL for each time point desired.

At the start of the measurement, doubly labeled Lphn3 GAIN domain was mixed with 2.5 equivalents of unlabeled GAIN domain in a final volume of 20 μL. For each time point, 2 μL of the GAIN domain mixture was sampled and mixed with 8 μL of HBS and 10 μL bead suspension. This mixture was incubated at room temperature while rotating end-over-end for 15 minutes. The beads were subsequently pulled down with a magnet, and 18 μL of the supernatant was aspirated to assess the protein content of the supernatant. The beads were resuspended in 16 μL HBS and stored at 4 ºC for elution prior to running the gel. Bead-bound protein was eluted by adding 6 μL of 4x LDS sample buffer (Thermo Fisher, NP0007) and heating the sample at 90 ºC for 5 minutes. The eluted sample was loaded into SDS-PAGE gels while pulling down the beads using a magnet. Cy3 fluorescence was assessed by imaging gels on a Typhoon FLA 9500 laser scanner (GE Healthcare Life Sciences). Gels were subsequently stained using Pierce Silver Stain Kit (Thermo Fisher, 24612) following manufacturers’ instructions.

Band intensities of silver-stained and fluorescence gel images were quantified using Fiji^47^. At each timepoint, an average value of the fraction of fluorescence in the bead fraction of protein relative to the total fluorescence in both bead and supernatant fractions was calculated. These mean fluorescence fraction values were fit to an exponential function *y=e*^*-bx*^, and the rate constant *b* was taken as the solution dissociation rate. 95% confidence intervals for the rate constants were calculated using the Jacobian from the exponential fits.

### GAIN domain magnetic tweezers dissociation assays

Magnetic tweezers were constructed and calibrated on a Nikon Ti-E microscope as previously described^48^. EGFP-YbbR-Lphn3 GAIN-HaloTag was biotinylated at the YbbR tag by reacting 4 μM protein with 10 mM MgCl_2_, 0.5 μM Sfp synthase, and 5 μM Biotin-PEG3-Coenzyme A (SiChem, SC-8618) at room temperature overnight. EGFP-YbbR-Lphn3 GAIN T855G-HaloTag was biotinylated similarly. PEG-passivated coverslips functionalized with HaloLigand were generated as previously described^28,29^. Coverwell perfusion chambers (Grace Bio-Labs, 622105) were placed on top of functionalized coverslips to generate several sample channels, and channels were blocked with 60 μL HBS with 1% casein (Millipore Sigma, C4765) overnight at 4 ºC in a humidity chamber. Dynabeads M-270 beads were prepared as follows: 1-10 μL of beads were washed three times with 300 volumes of HBS with 0.1% Tween, pulling down beads with a magnet and aspirating the supernatant each time. Beads were then resuspended in 30 volumes of HBS with 0.1% Tween and 1% casein and incubated rotating overnight at 4 ºC until use in experiments the next day.

On the day of the experiment, blocked coverslips were washed twice with 200 μL HBS. 60 μL of 10-30 nM EGFP-Biotin-Lphn3 GAIN-HaloTag diluted in HBS was added to channels and incubated at room temperature for 45 minutes to tether GAIN domains to the coverslip surface. Channels were treated with equal amounts of EGFP-Biotin-Lphn3 GAIN T855G-HaloTag as a non-cleavable control. As a negative control, channels were treated with either equal amounts of non-biotinylated EGFP-YbbR-Lphn3 GAIN-HaloTag or HBS. Channels were then washed three times with 200 μL HBS to remove any unreacted protein. Blocked beads were diluted 10x with HBS with 1% casein, for a final dilution of 1:300 from the original bead slurry. 60 μL of beads were added to the sample channels and incubated at room temperature for 15 minutes. The coverslip was then placed under the magnetic tweezers on the microscope and low (sub-pN) forces were applied to the sample channel for 3 minutes to dislodge non-adhered beads. Finally, the magnetic tweezers were brought to the height corresponding to the force of interest, and images from a 5×5 grid of fields of view were acquired at a rate of 1 image per field of view per minute over the course of 30-45 minutes.

### Bead dissociation assay data analysis

Bead counts were obtained from images by either manually counting beads or using the Particle Analysis plug-in in Fiji^47^ after applying a Gaussian filter to and thresholding acquired image stacks. For dissociation curves from non-cleavable Lphn3 T855G GAIN domain surfaces, bead counts were fit to the single exponential model

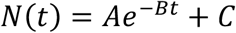

where *N* is the predicted bead count, *t* is time, *A* is an exponential prefactor, *B* is the estimated rate of bead dissociation, and *C* is a constant term that reflects beads that do not dissociate on the timescale observed. The extracted rate constants for non-specific dissociation across the forces measured were then fit to a linear model (Supp. Fig. 3) to estimate the average rate of non-specific dissociation as a function of force. For dissociation curves from wild-type Lphn3 GAIN domain surfaces, bead counts were fit to a biexponential model

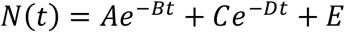

where *A* and *C* are exponential prefactors, *B* is constrained to be the predicted (slower) non-specific dissociation rate obtained from the Lphn3 T855G data, *D* is the dissociation rate reflecting the GAIN domain dissociation, and *E* is a constant term that reflects beads that do not dissociate on the timescale observed. The biexponential model was further subject to the constraint *A + C + E* = *N*_*0*_, where *N*_*0*_ is the bead count recorded at the start of the acquisition. The GAIN domain-specific dissociation rates are plotted as a function of force in Fig. 2C. We found that fits of the Lphn3 GAIN domain data to this constrained biexponential model were generally better than fits of the data to single exponential models (Supp. Table 1).

### GAIN domain tethered ligand assays

Purified, externally ligated GAIN domain was biotinylated by reacting 5 μM protein with 10 mM MgCl_2_, 0.5 μM Sfp synthase, and 5 μM Biotin-PEG3-Coenzyme A in HBS at 4 ºC overnight. The protein was then buffer-exchanged into HBS using a 7 kDa Zeba desalting column (Thermo Fisher, 89882) and stored at -80 ºC until use.

To generate glass surfaces functionalized with ELP-SpyCatcher, glass coverslips (Fisher Scientific, No. 12-544-14) were first cleaned and silanized as described previously^28,29^ to generate a nucleophilic, amine-functionalized substrate. Sulfosuccinimidyl 4-(N-maleimidomethyl)cyclohexane-1-carboxylate (Thermo Fisher, A39268) (Sulfo-SMCC) was diluted to 10 mM in 50 mM HEPES, pH 7.5, and a coverslip “sandwich” was formed by adding 90 μL of the Sulfo-SMCC solution between a pair of silanized coverslips. These coverslip sandwiches were incubated at room temperature in a humid environment for 1 hour to generate maleimide-functionalized coverslips. Coverslips were washed by dipping in a beaker of MilliQ water to remove excess Sulfo-SMCC, then gently dried under a stream of nitrogen. During this Sulfo-SMCC incubation, Cys-ELP-SpyCatcher was incubated with 5 mM tris(2-carboxyethyl)phosphine (TCEP) to reduce cysteines, then buffer exchanged into ELP conjugation buffer (50 mM phosphate, 50 mM NaCl, 10 mM EDTA, pH 7.2) using a PD MiniTrap G-25 (Cytiva Life Sciences, 28918007) desalting column. The final concentration of the reduced ELP solution was approximately 50 μM.

Coverslip sandwiches using the maleimide-functionalized coverslips were created with 90 μL of the ELP solution and incubated for 2 hours at room temperature in a humid environment. Coverslips were then washed by dipping in MilliQ water as described above and were sufficiently dry upon slow removal of the coverslips from the water due to the relatively hydrophobic surface properties of the ELP-functionalized surface. Finally, ELP-functionalized coverslip sandwiches were incubated with 100 μL of 10 mM L-cysteine in ELP conjugation buffer for 1 hour at room temperature in a humid environment to quench remaining free maleimides. These coverslips were then washed by dipping in water and were sufficiently dry for storage without additional drying steps. Coverslips were stored in vacuum-sealed bags at -20 ºC until further use.

3 channel Coverwell perfusion chambers (Grace Bio-Labs, 622103) were placed on top of the ELP-functionalized coverslip to create sample channels. Channels were blocked with 100 μL HBS with 1% casein in a humidity chamber. Dynabeads M-270 streptavidin beads were blocked by diluting with 20-300 volumes of HBS with 1% casein and 0.1% Tween as described above. Beads and sample channels were blocked either at 4 ºC overnight or at room temperature for a minimum of 1 hour. Following blocking, channels were washed with 200 μL HBS, then incubated with 110 μL biotinylated ligated GAIN domain protein at concentrations ranging from 2-50 nM for times ranging from 10 minutes to 1 hour at room temperature. Channels were washed with 2×200 μL HBS to remove unbound protein, after which blocked beads were directly added to the channel and incubated for 15-25 minutes at room temperature. Unbound beads were washed from the channel by slowly sweeping magnetic tweezers across the channel at a height corresponding to sub-pN forces for 20 seconds, then washing with 2×200 μL HBS. This washing was performed 3 times before the sample was measured.

Samples were illuminated from the objective side via a mint green LED (Thorlabs, MINTL5) reflected into the back focal plane of a 100x Plan-Apo TIRF objective using a polarizing beamsplitter (Edmund Optics). Reflected, cross-polarized light was transmitted through the beamsplitter and collected by a sCMOS camera (Hamamatsu Orca Flash v4). The *x-, y-*, and *z-*positions of the bead were recorded at 100 Hz from these images using a custom LabView script. The *x-* and *y-*positions were tracked using the 0^th^ order terms of the Fourier transform of the bead image. The *z-*position was computed using an empirical linear fit between the height and the phase of a high-amplitude frequency of the radial profile of the Fourier-transformed image. This fit was computed from a separately recorded look-up table of sample bead images at varying heights. Unbinding for individual molecules was probed by quickly increasing the force from sub-pN to the desired fixed force over the course of ∼2 seconds and tracking the *z-*position of the bead at the fixed force as an indicator of the binding state of the GAIN domain. Once the bead height exceeded a *z-*position greater than that corresponding to the unbound complex (i.e., the sum of ELP extension length and the GS linker length at the specified force), the force was decreased to sub-pN levels to allow the NTF and CTF of the complex to rebind. The complex was typically allowed to rebind for 120-180 seconds before the force was ramped back up to the desired force to probe unbinding.

### Tethered ligand assay data analysis and model fitting

Bead traces were fit using a custom Jupyter notebook to a piecewise function comprised of the following pieces:

1. A constant function capturing the initial tether height
2. An increasing linear function capturing the ELP extension from the initial force increase
3. A constant function reflecting the bead height immediately after ELP extension
4. A specified number of sequential constant functions, allowing for a variable number of steps, if needed
5. A decreasing linear function capturing the decrease in bead height as the force is reduced to sub-pN levels
6. A constant function capturing the tether height at the end of the acquisition under sub-pN forces

The analyzed traces were subjected to manual quality control checks and filtering to exclude recorded traces/dissociation events that were likely a result of processes not specific to the GAIN domain dissociation. These quality control criteria included verifying reasonable height extensions during the initial force ramp based on the contour length of the ELP, excluding traces that showed correlated *xy-*position changes during *z-*position jumps putatively corresponding to GS linker extension (reflecting the possible presence of multiple and/or non-specific tethers), excluding traces with implausible height changes after the initial extension based on a worm-like chain model^49^ of the GS linker, and excluding molecules with aberrant *xy-*fluctuations at low force that likely correspond to non-specific tethers.

For traces that met all quality control criteria and exhibited steplike behavior after the initial extension during the force ramp, the dissociation time was recorded as the time between the end of the force ramp and the step, during which the force was held constant. These dissociation times were compiled and plotted as binned lifetimes in Fig. 4A and as raw data in Supp. Fig. 4. We used maximum-likelihood estimation (MLE) to fit the lifetime data to one- and two-state slip bond models and generate values for model parameters that maximize the likelihood of observing the data by minimizing the negative log-likelihood function of each model. In the one-state slip bond model, the force-dependent rate of dissociation from bound state *1* to unbound state *0*, denoted *k*_*10*_*(F)*, follows Bell-Evans kinetics

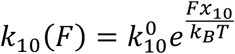

where 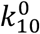 is the dissociation rate at zero force, *x*_*10*_ is a distance parameter for the transition, *k*_*B*_ is Boltzmann’s constant, and *T* is temperature. Then the likelihood of observing a specific dissociation time *t* at force *F* is given by

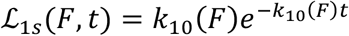

Thus, to find maximum-likelihood parameters 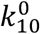 and *x*_*10*_, we minimize 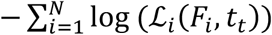 for the N=131 force-lifetime data points acquired in the tethered ligand assay.

For the two-state model, we adapt the two-state model described in Thomas et al.^50^, which specifies 4 force-dependent rates and 8 parameters, assuming a flux-balance initial condition. However, we found that a majority of the variation in the data could be captured with 4 parameters (i.e. increasing the number of parameters beyond 4 resulted in negligible marginal increases in the likelihood). In this model, the GAIN domain can interconvert between a strongly bound state *2* and weakly bound state *1* before dissociating to unbound state *0* in the manner depicted below

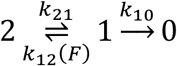

In this model, the transition rate between states *2* and *1* (*k*_*21*_) and the dissociation rate of the complex (*k*_*10*_) are considered force insensitive, whereas the reverse transition from state *1* to state *2* is force sensitive according to Bell-Evans kinetics such that

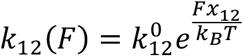

and *x*_*12*_ is fixed to be negative such that the rate of transition to strongly bound state *2* decreases as force is increased and the overall bond lifetime decreases with force as expected for a slip bond. Following the convention in Thomas et al., we can define

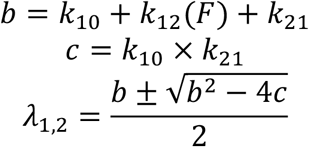

We further define *C*_*1*_ and *C*_*2*_ as

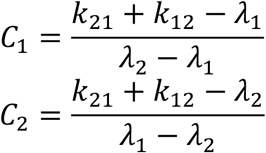

In this model, the likelihood of observing a specific dissociation time *t* at force *F* is given by

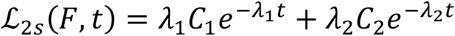

We similarly minimize the summed negative log-likelihood functions across all points in the data set to find the maximum-likelihood parameters for this two-state model. Optimization was performed using the Matlab implementation of the genetic algorithm, with 100 epochs. Mean dissociation times at specific forces predicted from one- and two-state models were calculated as 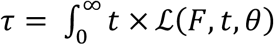 *dt* where *θ* denotes the MLE fit parameters. For the two-state model, 95% confidence intervals on the parameters and the dissociation times at the force range covered by the data were computed as the 2.5 and 97.5 percentiles of fit parameters and predicted dissociation times from fits to 1000 synthetic datasets, sampled with replacement. For bootstrapped fits to synthetic datasets, 50 epochs of the genetic algorithm were used.

## Notes

### Competing Interest Statement

The authors have declared no competing interest.

