## Supplementary Info for "Piconewton forces mediate GAIN domain dissociation of the latrophilin-3 adhesion GPCR"

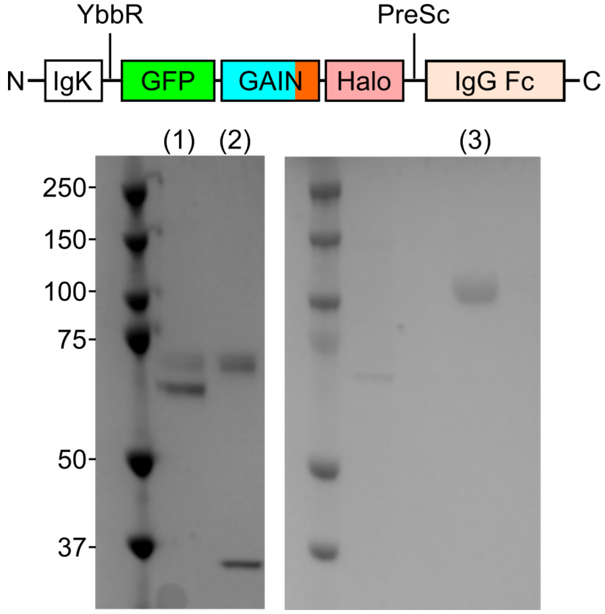


**Supplementary Figure 1. Expression of Lphn3 GAIN domain construct for magnetic tweezers bead dissociation assay**

Secreted Lphn3 fusion protein *(top)* was expressed in Expi293 HEK cells and purified from the supernatant using a Protein A column. Reducing gels depict Lphn3 GAIN domain fusion protein prior to *(1)* and after *(2)* on-column cleavage using PreScission protease. A fusion protein with the Lphn3 T855G mutant GAIN domain was expressed and purified using the same strategy and runs at a molecular weight consistent with the sum of the NTF and CTF, depicted after PreScission cleavage in *(3).*

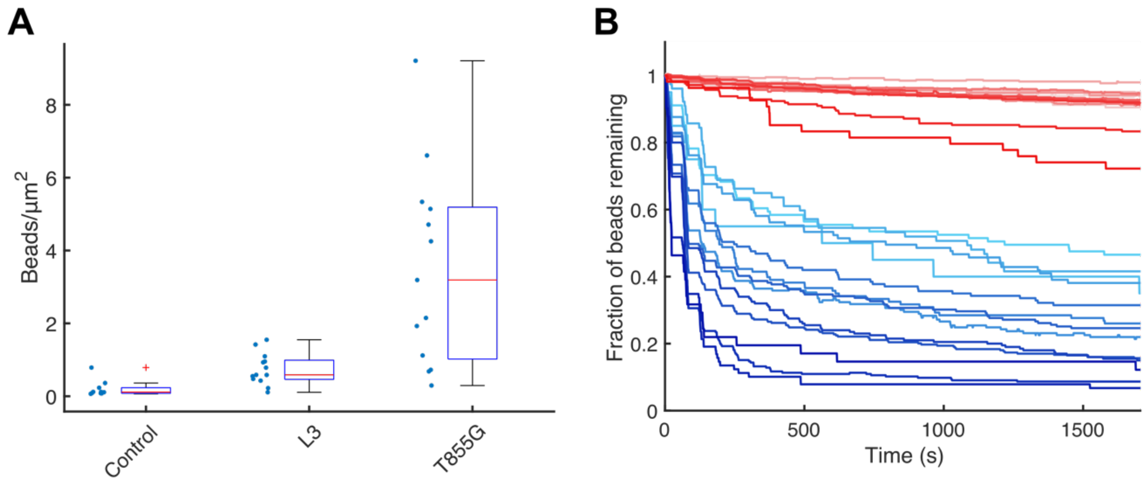


**Supplementary Figure 2. Bead counts and all traces for Lphn3 GAIN domain bead dissociation assay**

**(A)** Bead densities on control surfaces (either no protein or non-biotinylated protein, both shown combined), Lphn3 GAIN functionalized surfaces (L3), and Lphn3 T855G GAIN functionalized surfaces for bead dissociation assay. **(B)** Fraction of beads remaining over time for Lphn3 GAIN surfaces (blue) and Lphn3 T855G GAIN surfaces (red), where darker shade indicates higher force ranging from 1.7 pN to 9.0 pN. First 30 minutes of acquisitions are shown for comparison.

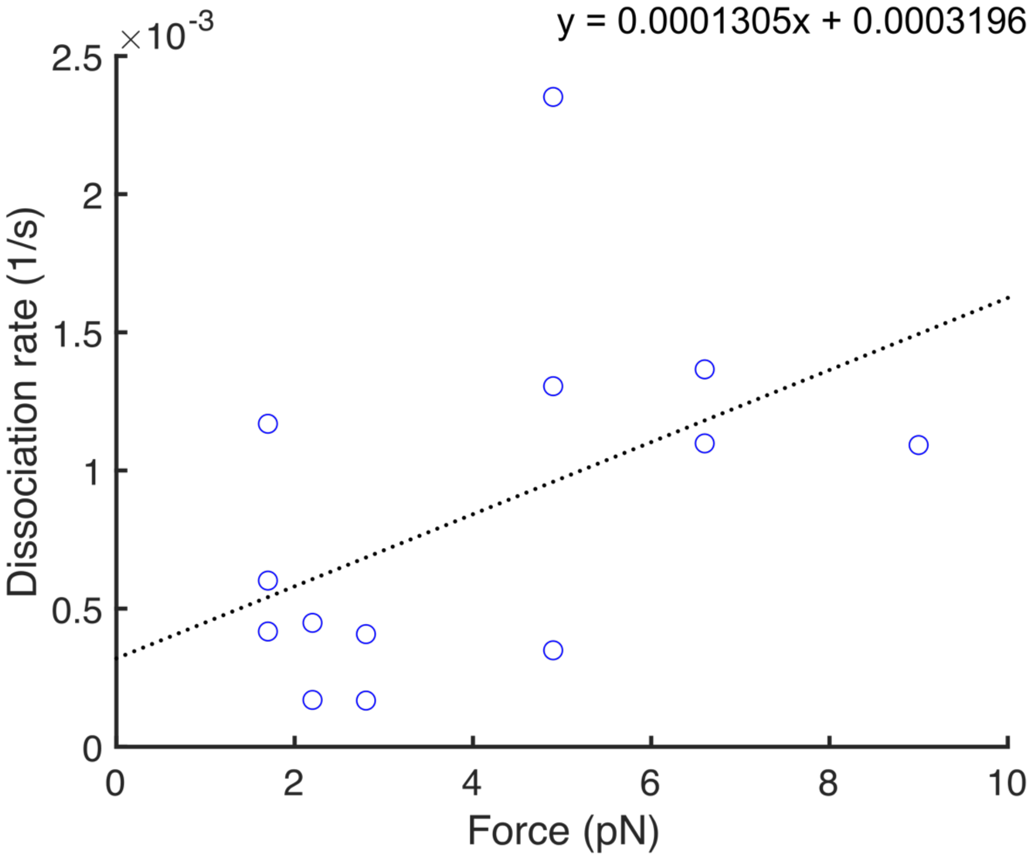


**Supplementary Figure 3. Non-specific rates of bead dissociation in Lphn3 bead dissociation assay**

Non-specific rates of bead dissociation were estimated by measuring bead dissociation from Lphn3 T855G GAIN domain surfaces and fitting these data to a single exponential plus constant ($Ae^{-Bx} + C$). Rates from these fits are plotted as blue dots above and fit to a linear fit to interpolate non-specific dissociation rates at a given force.

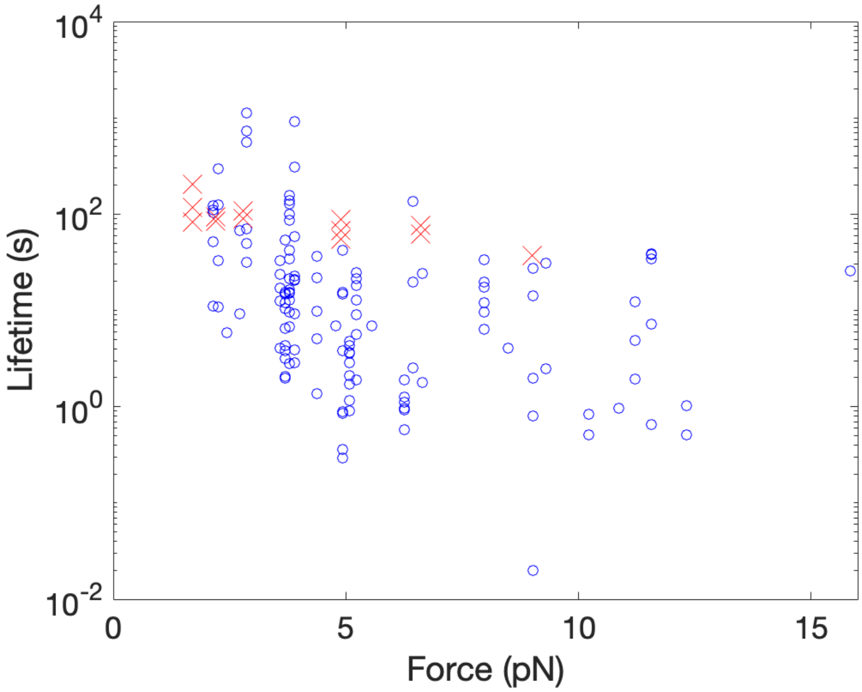


**Supplementary Figure 4. Lifetimes from tethered ligand assay and bulk dissociation assay**

Lifetimes measured from tethered ligand assay are depicted in blue circles, with each circle representing one bond rupture event (N = 131 events from 19 molecules). Calculated lifetimes from bulk dissociation assay (Fig. 2, Supp. Table 1) are overlaid in red for comparison.

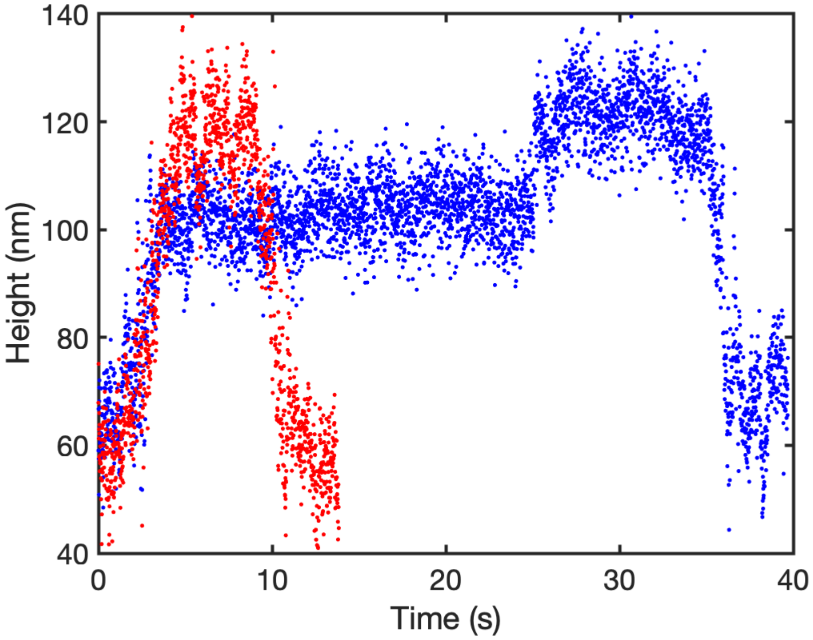


**Supplementary Figure 5. Sample data for tethered ligand GAIN molecule that has not yet rebound**

Comparison of bead height data for a trace with a distinct second step *(blue)*, representing a bound tethered ligand GAIN domain that unbinds, with a trace from the same molecule that reaches the unbound bead height immediately after the force is increased to the desired force *(red)*. Lack of a distinct step and direct extension to full unbound bead height suggest that the molecule has not yet rebound during the waiting period between force application.

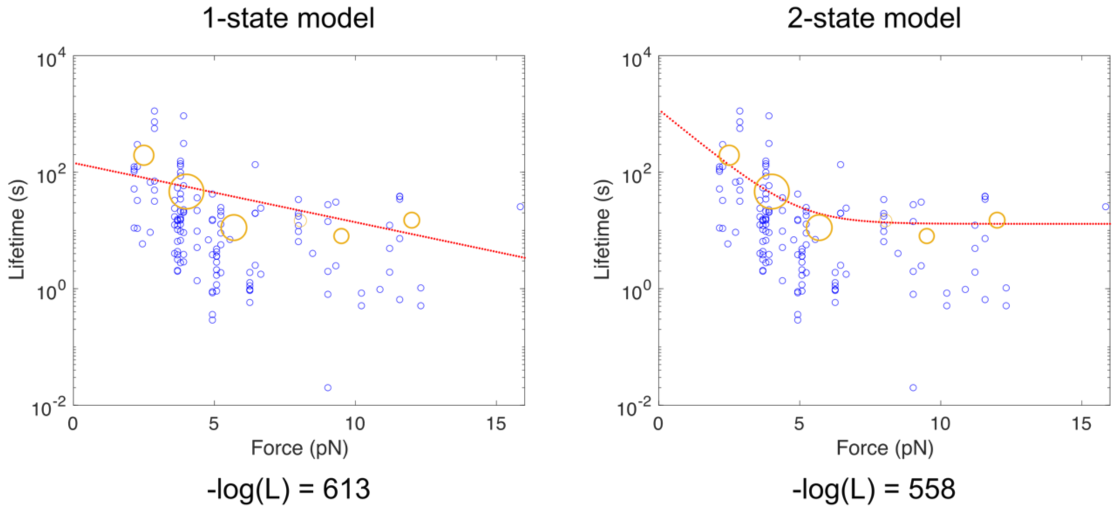


**Supplementary Figure 6. Comparison of 1- and 2-state fits to tethered ligand data**

Maximum-likelihood estimate fits of tethered ligand data to 1- and 2-state dissociation models indicate that a 2-state model is more likely based on the log-likelihood values of fits. Individual force-lifetime data points are plotted as blue circles, with larger orange circles representing the average data across 2 pN wide bins (except for the highest force bin, which also includes the single data point at ~16 pN). Size of orange circle scales with number of data points in the bin. Predicted lifetimes from the two fits are plotted as red points.

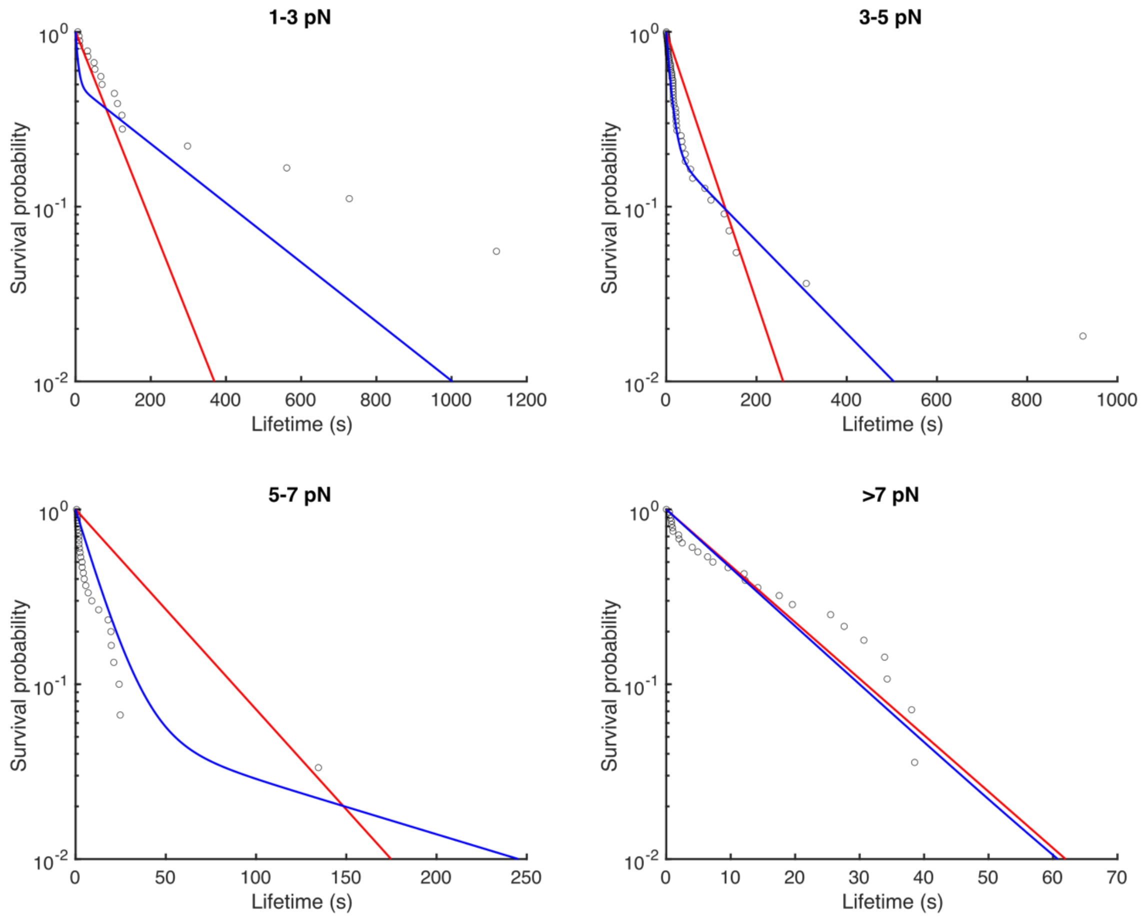


**Supplementary Figure 7. Survival probability curves for binned Lphn3 tethered ligand dissociation lifetimes**

Survival probability curves are plotted for various force bins for Lphn3 tethered ligand dissociation lifetimes, with predicted survival probabilities from one-state *(red)* and two-state *(blue)* models overlaid. Data are better fit by a two-state model.

**Supplementary Table 1. Single- and double-exponential fit parameters for Lphn3 GAIN domain bead dissociation data**

| | Single Exponential | | Double Exponential | | |
|---|---|---|---|---|---|
| **Force (pN)** | **Rate ($s^{-1}$)** | **Adj $R^2$** | **Fast Rate ($s^{-1}$)** | **Slow Rate ($s^{-1}$, fixed)** | **Adj $R^2$** |
| 1.7 | 0.00206 | 0.921 | 0.00854 | 0.000541 | 0.988 |
| 1.7 | 0.00192 | 0.965 | 0.00485 | 0.000541 | 0.977 |
| 1.7 | 0.00147 | 0.900 | 0.01199 | 0.000541 | 0.966 |
| 2.2 | 0.00129 | 0.951 | 0.01058 | 0.000607 | 0.993 |
| 2.2 | 0.00300 | 0.895 | 0.01193 | 0.000607 | 0.985 |
| 2.8 | 0.00484 | 0.966 | 0.00928 | 0.000685 | 0.990 |
| 2.8 | 0.00261 | 0.919 | 0.01112 | 0.000685 | 0.990 |
| 4.9 | 0.00546 | 0.898 | 0.01817 | 0.000959 | 0.987 |
| 4.9 | 0.00804 | 0.938 | 0.01496 | 0.000959 | 0.985 |
| 4.9 | 0.00607 | 0.946 | 0.01125 | 0.000959 | 0.982 |
| 6.6 | 0.01152 | 0.973 | 0.01310 | 0.001181 | 0.979 |
| 6.6 | 0.01128 | 0.975 | 0.01607 | 0.001181 | 0.955 |
| 9.0 | 0.01454 | 0.922 | 0.02706 | 0.001494 | 0.953 |

**Supplementary Note 1. Estimating on-rate and $K_d$ of Lphn3 NTF-CTF interaction**

The data in Fig. 4 and Supp. Fig. 4 depict 131 dissociation events from 19 distinct Lphn3 GAIN domain molecules. However, not all traces acquired for these molecules yielded unbinding events, as illustrated in Supp. Fig. 5. In total, we acquired 636 traces from these 19 molecules during which GAIN domains spent a total of 72756 s under low force (< 1 pN) to allow for rebinding of the GAIN domain NTF and CTF. Thus, the average "rebinding time" for the NTF-CTF in the tethered ligand assay is 72756/131 = 555 s.

We can estimate the effective concentration of NTF and CTF by taking the approximate volume of the tethered ligand protein construct—and more specifically the GS linker connecting the NTF and CTF—to be the volume in which the NTF and CTF reside. This volume is on the order of 100 $nm^3$, which approximately corresponds to a sphere of radius 2-3 nm. A density of 1 molecule per 100 $nm^3$ gives a concentration of 16.6 mM. Using this concentration, we compute a characteristic on-rate $k_{on}$

$$k_{on} = \frac{1}{(0.0166\,M)(555\,s)} = 0.11 M^{-1}s^{-1}$$

From solution dissociation measurements in HEPES-buffered saline (Fig. 1), we estimate the off-rate $k_{off}$ of the NTF-CTF interaction to be 0.0011 $hr^{-1}$ = $3.1 \times 10^{-7}$ $s^{-1}$. Thus, the binding affinity $K_d$ of the Lphn3 NTF and CTF can be estimated to be

$$K_d = \frac{k_{off}}{k_{on}} = \frac{3.1 \times 10^{-7} s^{-1}}{0.11\, M^{-1}s^{-1}} = 2.8\,\mu M$$
